# Destructive and constructive cheater suppression through quorum sensing

**DOI:** 10.1101/2021.06.16.448736

**Authors:** Alexander S. Moffett, Peter J. Thomas, Michael Hinczewski, Andrew W. Eckford

## Abstract

The evolutionary consequences of quorum sensing in regulating bacterial cooperation are not fully understood. In this study, we reveal unexpected consequences of regulating public good production through quorum sensing on bacterial population dynamics, showing that quorum sensing can be a collectively harmful alternative to unregulated production. We analyze a birth-death model of bacterial population dynamics accounting for public good production and the presence of non-producing cheaters. Our model demonstrates that when demographic noise is a factor, the consequences of controlling public good production according to quorum sensing depend on the cost of public good production and the presence of non-public fitness benefits. When public good production is inexpensive, quorum sensing is a destructive alternative to unconditional production, in terms of the mean population extinction time. When costs are higher, quorum sensing becomes a constructive strategy for the producing strain, both stabilizing cooperation and decreasing the risk of population extinction.

## 1 Introduction

Cooperative behavior is widespread in bacteria [1], including coordinated swarming and public good production. These behaviors are often regulated through quorum sensing (QS) [2], in which individual bacteria produce and export small molecules called autoinducers (AI). When AI molecules accumulate to a sufficiently high concentration in the environment, and consequently within the bacteria producing them, they activate operons controlling the expression of genes critical for cooperation. While the biochemistry of some QS systems is well understood [2], there are many proposed biological functions of QS which have been the subject of debate [3, 4, 5, 6, 7].

Evolutionary questions concerning QS function have largely focused on the ability of QS-controlled cooperation to combat invasion by non-cooperating cheaters in bacterial populations [8, 9]. The question of how cooperation can evolve and be maintained in populations is a general problem in evolutionary biology [10], and the role of QS in the evolution of bacterial cooperation is of great interest in understanding bacterial social interactions. A number of possible resolutions to the problem of social cheaters in bacterial public good production include punishment of cheaters [11], dispersal into subpopulations [12], and the use of QS to regulate cooperation [1, 13, 6]. Regulation of public good production through QS has been shown to reduce the ability of cheaters to invade a population of producers [14].

The role of QS in maximizing population growth in the absence of cheaters has also been investigated as a rationale for QS-control of public good production. Because public goods produced by bacteria can have density-dependent fitness benefits [15], regulation of public good production based on population density can be seen as an optimal control solution to maintaining a maximal population size balancing metabolic costs and benefits [16, 17]. By maintaining a maximal population size, a population also maximizes its mean time to extinction, which is especially important with the possibility of unpredictable environmental changes.

Considering the threat of both cheater invasion and the onset of harmful environmental conditions that could lead to population extinction, there is a tension in the degree to which public good production is regulated. Unconditional public good production appears to be a self-defeating strategy, where non-producing cheaters will arise by mutation and reap all the benefits of public goods without experiencing any of the associated production costs. At the other extreme, a strategy where no public goods are produced (which could be a result of a successful cheater invasion of a cooperating population) should be vulnerable to extinction when the public goods are essential for growth or reducing the likelihood of death. The idea that quorum sensing is a moderate strategy between these two extremes has been explored previously [18]. However, to our knowledge, a full analysis of this tension explicitly considering both cheater fixation probability and mean population extinction time has not been carried out. What are the effects of QS-mediated regulation of public good production in terms of cheater suppression and overall population robustness? Does QS always protect against cheaters while also increasing long-term viability of the population?

In this work, we explore the effects of QS on cheater fixation and mean population extinction time in simple birth-death models of mixed producer-cheater populations. We compare the QS strategy of public good production with an “always on” (AO) strategy, meaning that each producer cell unconditionally produces public goods at the maximum rate. Our models reflect bacterial populations where demographic stochasticity is an important factor in population dynamics. The role of demographic stochasticity in bacterial populations has been explored in past work [19, 20], and is particularly important when populations are divided into subpopulations, as is the case with *Pseudomonas aeruginosa* lung infections [21] for example. Additionally, Joshi and Guttal recognized demographic noise as a possible factor in the evolution of QS in populations of fixed size [22]. We use the QS system controlling production of proteases by *P. aeruginosa* as inspiration for our model, where public goods increase the growth rate of all individuals while leaving death rates unaffected.

We find that while QS decreases cheater fixation probability for all examined public good costs and constitutive growth rates (growth rate in the absence of public goods), the population mean extinction time is only increased by QS for a well-defined set of cost-growth rate pairs. The cases where QS decreases mean extinction time as compared with an unconditional AO strategy are analogous to spiteful behavior [23, 24], where QS reduces the relative fitness of the cheater strain while the entire population of both producers and cheaters is made more vulnerable to extinction. We call this apparently “spiteful” behavior destructive cheater suppression. Only when both the cost of public good production and the constitutive growth rate are large enough is QS a constructive strategy for the producers, in that both cheater fixation probability is reduced and mean extinction time is increased. We discuss the relationship between our definitions of destructive and constructive cheater suppression with the well-studied notions of evolutionary spite and selfishness in the discussion section.

## 2 Results

We first briefly introduce the notation used throughout the remainder of this article. The number of producers in the population is written as *n* while the number of cheaters is *m*. The cost of public good production is *c* and the per-capita constitutive growth rate (in the absence of public goods) is λ_0_. The per-capita death rate for all individuals is *μ*_0_. The birth rates given *n* and *m* are 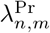 and 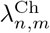 while the death rates are 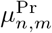 and 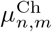, for the producers and cheaters, respectively. We write the cheater fixation probability as *π*^Ch^, the mean time to cheater fixation as *τ*^Ch^, and the mean time to population extinction as 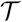. The model is fully described in the Materials and Methods section.

The phase diagram in Fig. 1B shows the fraction of (*n, m*) pairs in the set 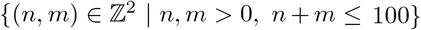 in which the mean extinction time of a population regulating public good production through QS is larger than that using the AO strategy, as a function of the constitutive per capita growth rate, λ_0_ ≥ 0, and the cost of public good production, 0 ≤ *c* ≤ 1, representing the fraction of growth rate benefit provided by public goods alone. We first examine two points of interest on this phase diagram, *c* = 0.1, λ_0_ = 0 and *c* = 0.15, λ_0_ = 0.2 as marked in Fig. 1B, and then discuss the general features of the phase diagram.

**Figure 1:**
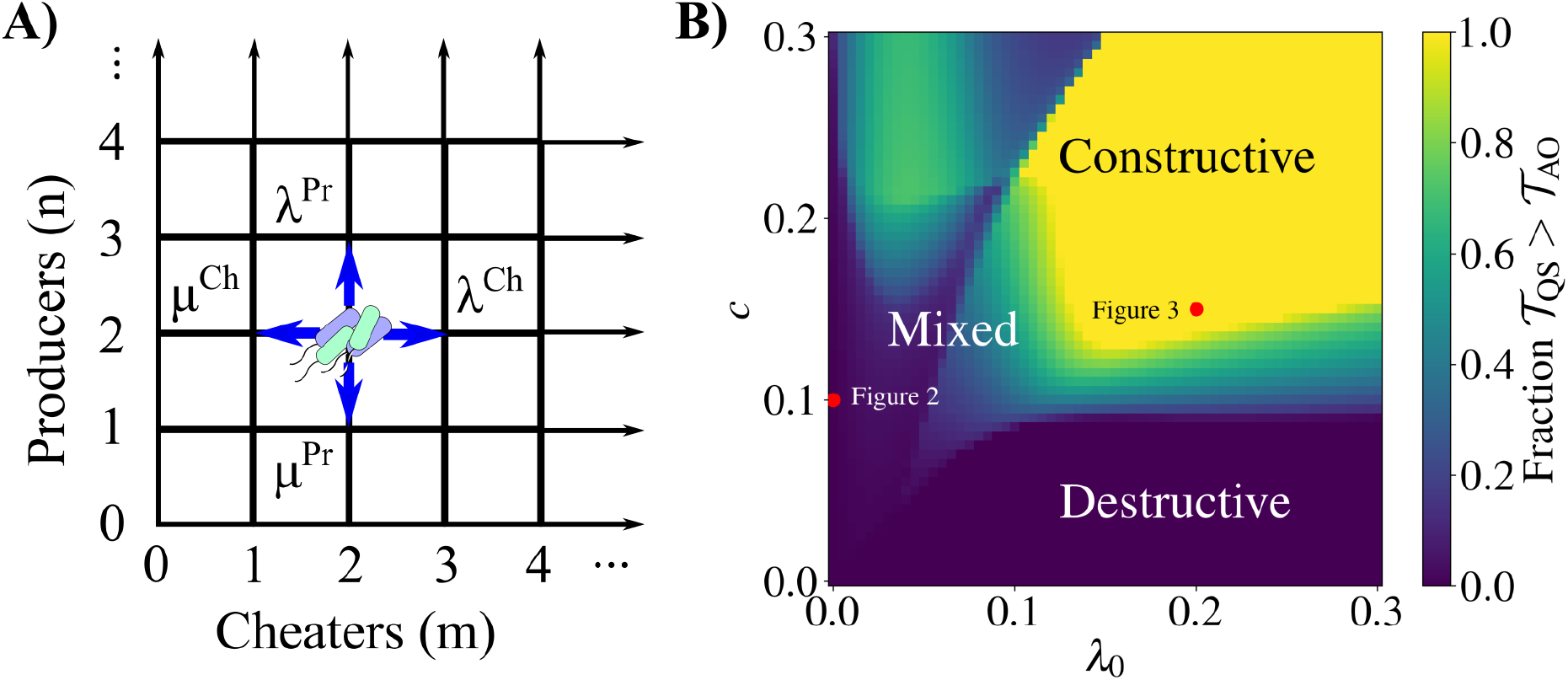
Overview of birth-death model and main results. A) Diagram of the two-dimensional birth-death process describing the dynamics of a bacterial population consisting of public good producers and cheaters. As an example, the population is shown with two producers and two cheaters. All possible subsequent states of the population are indicated, where a single producer or cheater can either arise through binary fission (with state-dependent rates λ^Pr^ and λ^Ch^, respectively) or can die (with state-dependent rates *μ*^Pr^ and *μ*^Ch^). See Eqs. 1–4 for the full forms of the birth and death rates. B) Phase diagram describing the fitness gains of quorum sensing (QS) over always on (AO) producers, as a function of constitutive growth rate (λ_0_) and public good production cost (*c*). For each pair (λ_0_, *c*) we calculated the mean extinction time for all (*n, m*) pairs satisfying *n* ≥ 0, *m* ≥ 0, and *n* + *m* ≤ 100 with the QS and AO strategies. The reported number is the fraction of these (*n, m*) pairs for which the mean extinction time for QS is greater than for AO 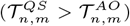. In the bottom right-hand region (dark blue) regulating public good production through QS decreases mean extinction time for all (*n, m*) pairs, meaning that the AO strategy decreases the risk of population extinction. Here, QS is a destructive strategy. The upper region (yellow) is where QS increases the mean extinction time for all (*n, m*), a constructive strategy for the producers. The region labeled “mixed” indicates that QS increases mean extinction time for some (*n, m*) pairs while decreasing it for others.

### 2.1 Quorum sensing is a destructive strategy in the absence of alternative nutrition sources

In agreement with previous work [14, 25], we find that in the absence of alternate sources of nutrition (λ_0_ = 0), QS reduces the probability of cheater fixation for a wide range of starting population compositions (Fig. 2). We calculate the probability of cheaters fixing (Eq. 9) in a small population with starting compositions satisfying 0 < *n*, 0 < *m*, and *n* + *m* ≤ 100. We compare cases where no public good is produced (NP), where public good production is QS-controlled, and where public good production is always on (AO). Because there is no constitutive growth rate (λ_0_ = 0), if the NP strategy is used there is no (*n, m*) pair for which the net population growth rate is non-negative. On the other hand, for the QS and AO populations, the white lines in Fig. 2 represent the total population zero expected net growth contour. Because these lines represent the zero expected net growth rate contours of the entire population, they are not fixed points of the underlying deterministic behavior of the system, which only exist at the origin and at the intersection of the zero expected net growth contours with the producer axis. Thus, there can not be a long-lived population with a non-zero number of cheaters.

**Figure 2:**
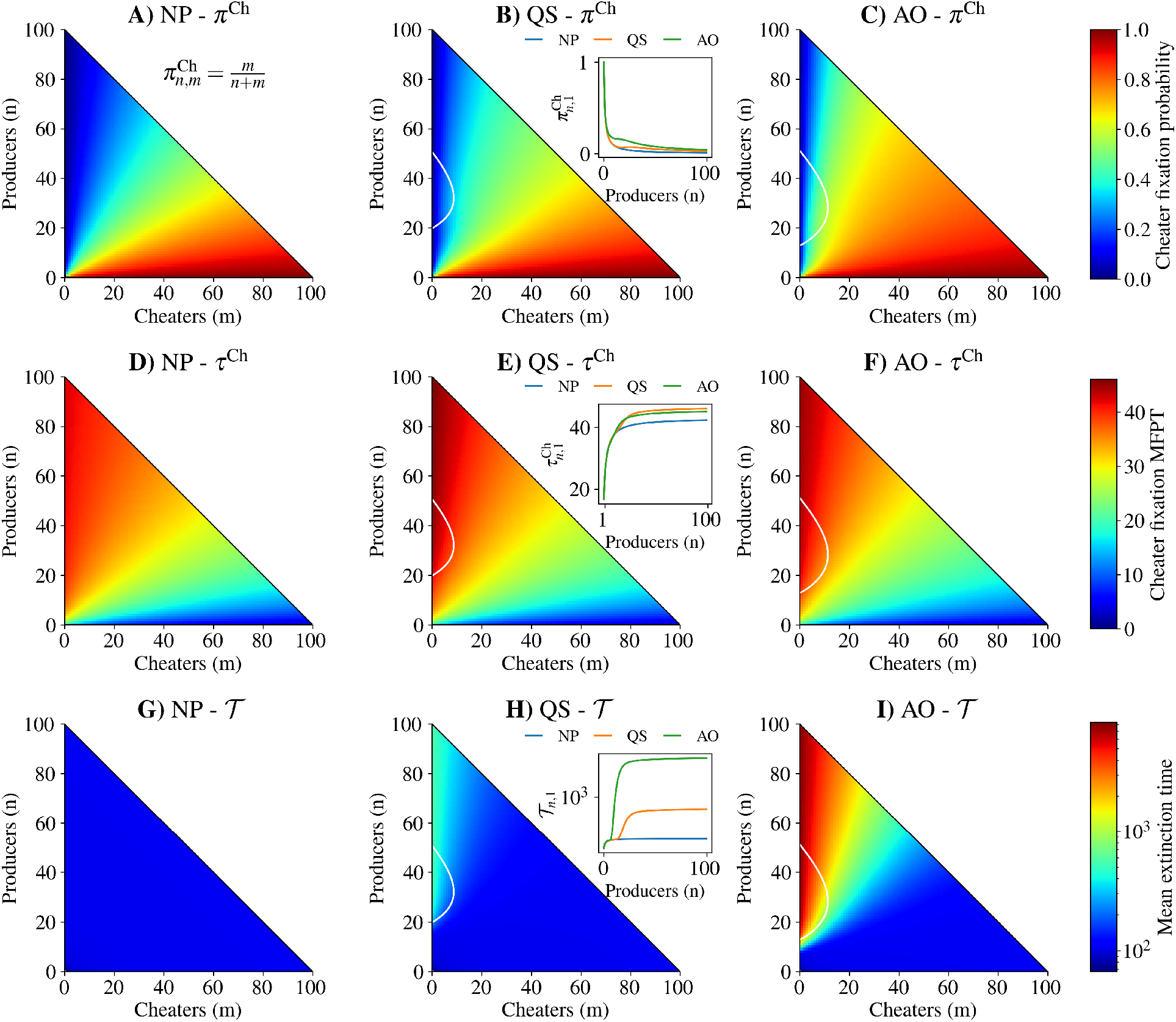
Quorum sensing is a destructive strategy for producers when no alternate energy source is present (λ_0_ = 0) and when public good cost is moderate (*c* = 0.1). Row 1: Cheater fixation probability from an initial population of *n* producers and *m* cheaters for A) no production (NP), B) quorum sensing (QS), and C) always on (AO) strategies. Row 2: Conditional mean first passage time to cheater fixation from initial population structure for D) no production, E) quorum sensing, and F) always on strategies. Row 3: Mean extinction time from initial population structure for G) no production, H) quorum sensing, and I) always on strategies. White traces: zero expected net total population growth contours. Inset plots in B), E), and H) show cheater fixation probability, cheater mean first passage time to fixation, and mean extinction time, respectively, for an initial population with one cheater and *n* producers. See Table 1 for parameter values.

With a moderate cost to public good production (10% of the maximum growth rate benefit imparted by public goods), a QS strategy (Fig. 2B) increases cheater fixation probability as compared with an NP strategy (Fig. 2A) while decreasing cheater fixation probability as compared with an AO strategy (Fig. 2C). This can be clearly seen in an invasion scenario with a single cheater present in the population (Fig. 2C inset). For any public good production strategy, without any elaborate solutions such as policing [26] it is unavoidable that cheaters will become more likely to fix in a population. However, QS mitigates this possibility as compared with an AO strategy. Both the QS and AO strategies (Fig. 2E-F) increase the mean time to extinction as compared with the NP strategy (Fig. 2D). Our results match well with simulations (Figs. S1 & S2).

In order to explore the fitness advantages of public good production to the total population, we calculated the mean time to extinction of the population given an initial composition of producers and cheaters. The mean extinction time has been proposed as a general measure of fitness [27] and has the advantage over geometric mean growth rate [28] that it can be applied to populations with limited growth. While QS reduces the probability of cheater fixation as compared to AO, the mean extinction time with QS (Fig. 2H) is greatly reduced as compared to AO (Fig. 2I). This suggests that with no alternate sources of nutrition, QS increases the relative fitness of producers while decreasing the overall population fitness as compared with AO. This scenario is reminiscent of evolutionary spite [23, 24] which we call destructive cheater suppression. For most, but not all, (*n, m*) pairs, the mean extinction time for QS is larger than for AO. This places the system in the “mixed” region of the phase diagram (Fig. 1B).

The qualitative differences in cheater fixation probabilities and mean extinction times between QS and AO were preserved for systems with carrying capacities of 200 (Fig. S3), twice the size considered in Figs. 2 (see Tables 1 & 2 for the parameters used). While this does not guarantee that the destructive nature of QS in these conditions is independent of populations size, it does demonstrate insensitivity to the overall population size. Given that our model is not parameterized by experimental results, the population sizes we consider do not directly correspond to population sizes in real bacterial colonies. Instead, our model aims to capture key aspects of bacterial population dynamics when demographic noise plays a role, as discussed in the Introduction. The degree of demographic noise is related to the size of a population through the law of large numbers, where the standard deviation in the population size over the mean number of individuals is inversely proportional to the square root of the population size. For this reason we focus on small populations, where demographic noise is relatively large.

**Table 1:** Model parameters used in main text figures.

| Parameter (Unit) | Figure 1 | Figure 2 | Figure 3 | Figure 4 |
| --- | --- | --- | --- | --- |
| $\lambda_0$ (1/time) | Variable | 0.0 | 0.2 | 0.2 |
| $\mu_0$ (1/time) | 0.015 | 0.015 | 0.015 | 0.015 |
| $g$ (1/time) | 1.0 | 1.0 | 1.0 | 1.0 |
| $c$ (1/time) | Variable | 0.1 | 0.15 | 0.15 |
| $K_g$ (unitless) | 20.0 | 20.0 | 20.0 | 20.0 |
| $h_g$ (unitless) | 2.0 | 2.0 | 2.0 | 2.0 |
| $K_a$ (unitless) | 15.0 | 15.0 | 15.0 | 15.0 |
| $h_a$ (unitless) | 4.0 | 4.0 | 4.0 | 4.0 |

**Table 2:**
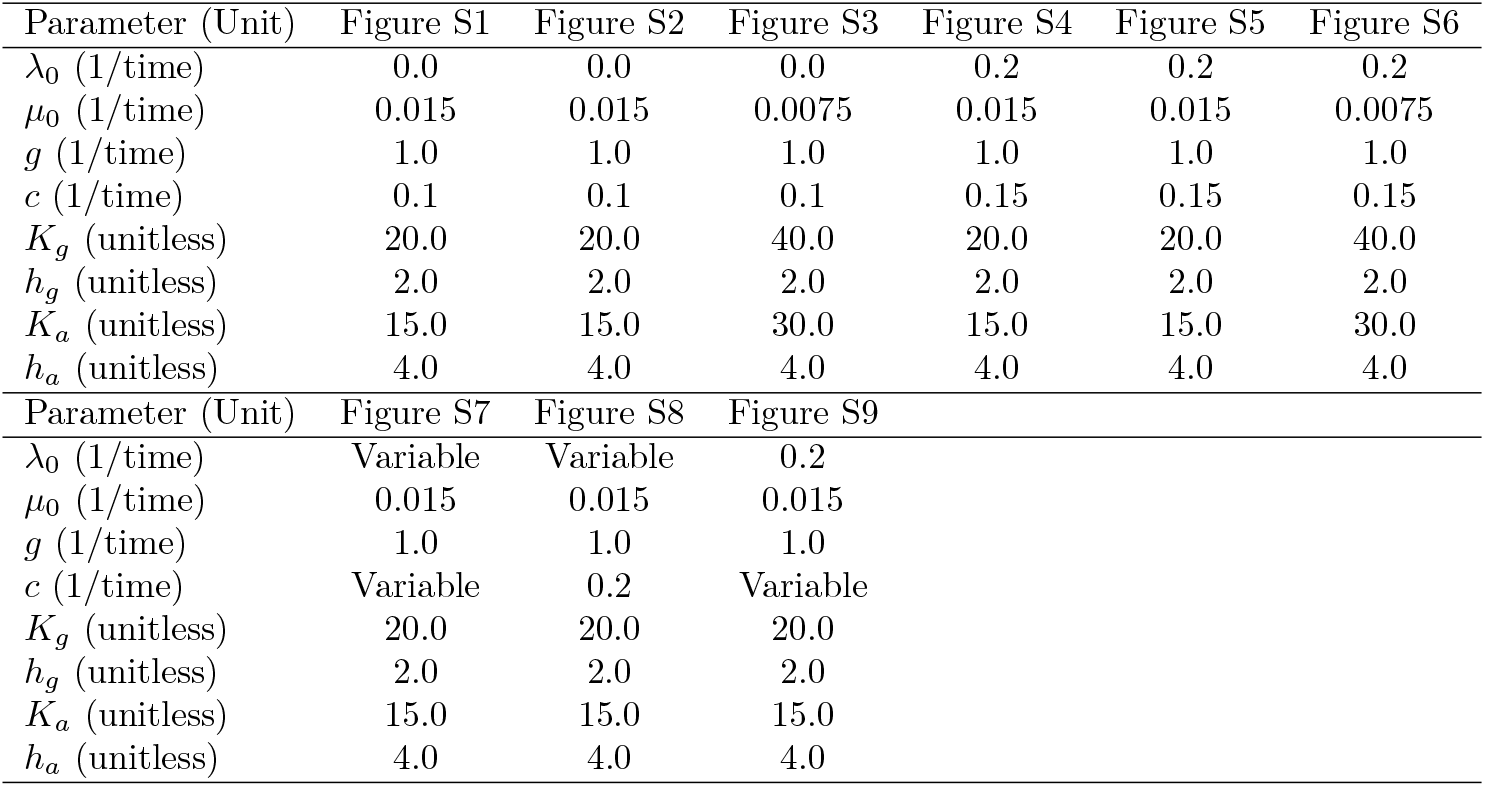
Model parameters used in supplemental figures.

### 2.2 Alternative sources of nutrition enable constructive suppression through QS

Bacterial populations need not rely solely on the growth benefits of a public good. In some cases they may use alternative sources of nutrition which are not directly influenced by public good production. We consider the case in which the constitutive growth rate for individuals with zero public goods present is non-zero but small (20% of the maximal growth rate benefit provided by public goods alone). This non-zero constitutive growth rate allows for a small carrying capacity to appear in the NP case, indicated by the white line in the leftmost column of Fig. 3. With the QS and AO strategies, the zero expected net growth contour reflects the NP carrying capacity when the number of producers is low, but increases markedly with more producers. Additionally, more cheaters can be accommodated with more producers present because of the public nature of the fitness benefits provided by producers. As in Fig. 2, the only fixed points of the underlying deterministic dynamics are the origin and the intersections of the zero expected net growth contour with the axes.

**Figure 3:**
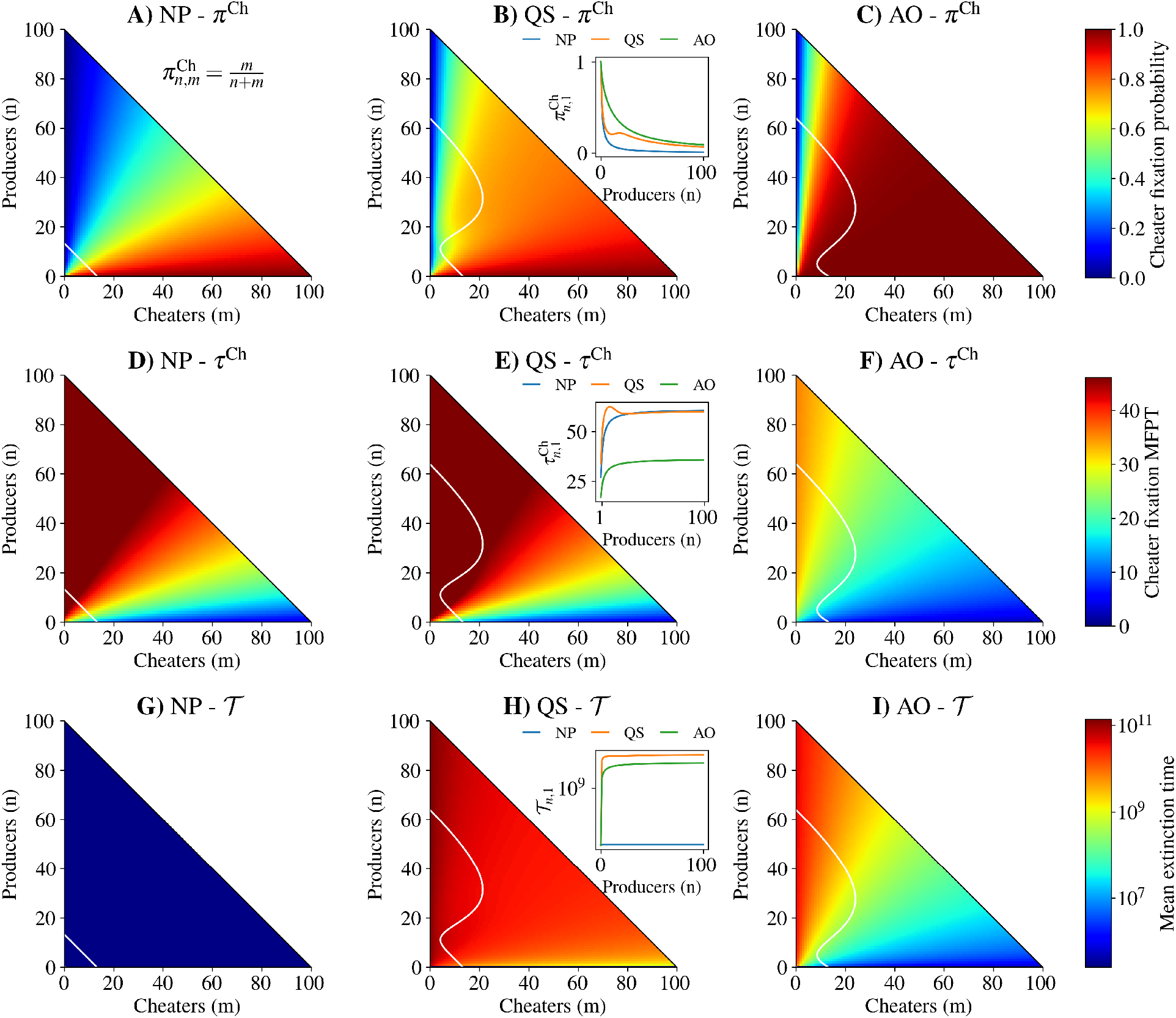
Quorum sensing is a constructive strategy for producers when an alternate energy source is present (λ_0_ = 0.2) and when public good cost is moderate (*c* = 0.15). Row 1: Cheater fixation probability from an initial population of *n* producers and *m* cheaters for A) no production (NP), B) quorum sensing (QS), and C) always on strategies (AO). Row 2: Conditional mean first passage time to cheater fixation from initial population structure for D) no production, E) quorum sensing, and F) always on strategies. Row 3: Mean extinction time from initial population structure for G) no production, H) quorum sensing, and I) always on strategies. White traces: zero expected net total population growth contour. Inset plots in B), E), and H) show cheater fixation probability, cheater mean first passage time to fixation, and mean extinction time, respectively, for an initial population with a single cheater and *n* producers. See Table 1 for parameter values.

As in the case of zero constitutive growth, QS (Fig. 3B) decreases the probability of cheater fixation over AO (Fig. 3C) for a wide range of starting population compositions, while increasing the fixation probability compared to NP (Fig. 3A). The improvement of QS over AO can be clearly seen in the cheater invasion scenario in the inset of Fig. 3B. However, with non-zero constitutive growth the mean time to cheater fixation for NP (Fig. 4D) and QS (Fig. 3E) are comparable, while cheaters fix more quickly for AO (Fig. 3F). Again, our results match well with simulations (Figs. S4 & S5).

**Figure 4:**
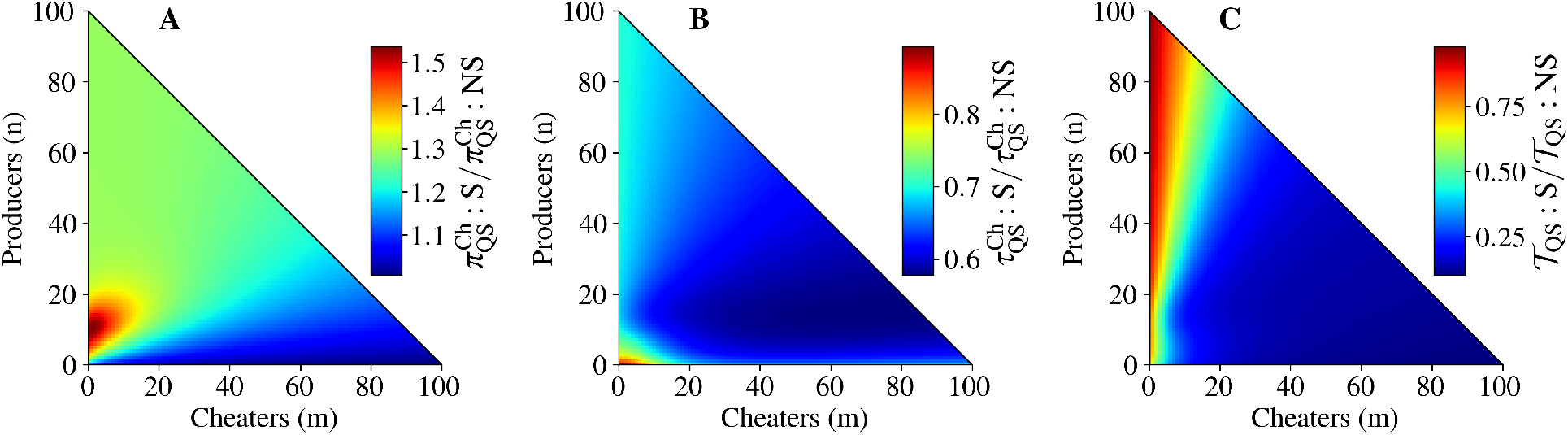
Cheater autoinducer production is destructive in the presence of an alternative source of nutrition (λ_0_ = 0.2). The ratio of A) cheater fixation probability, B) cheater mean fixation time, and C) mean extinction time between QS populations where cheaters signal (indicated by *S*) to where cheaters do not signal (indicated by *NS*). The non-signaling results used in the denominators are the same as those in Fig. 3. See Table 1 for parameter values.

The presence of an alternative source of nutrition renders QS a beneficial strategy for producers (Fig. 3) at a moderate public good production cost (15%). QS increases the mean extinction time of the population for all starting population compositions with at least one cheater and one producer over AO (Fig. 3H-I). From comparison with (Fig. 2), it appears that the degree to which growth depends on alternative nutrition sources influences whether QS at moderate costs is destructive or constructive.

As when there is no constitutive growth, the qualitative differences between QS and AO were preserved for systems with carrying capacities of 200 as shown in Fig. S6 (see Tables 1 & 2 for parameters).

### 2.3 Autoinducer production by cheaters is a destructive strategy

To this point, we have assumed that cheaters produce neither public goods nor autoinducer. What are the consequences of cheaters producing autoinducer while still not producing public goods? We performed the same analysis as in the previous sections using the growth rates in Eqs. 5 & 6 which reflect equal autoinducer production by producers and cheaters. We compared these results to those presented in Fig. 3, where an alternative nutrition source exists and cheaters do not produce autoinducer. Cheater signaling increases the probability of cheater fixation while decreasing the mean time to cheater fixation and the mean extinction time (Fig. 4). From the perspective of the cheater population, autoinducer production is a destructive strategy. Even though cheater signaling decreases mean extinction time, this result suggests that signaling cheaters are more likely to be observed in nature than non-signaling cheaters, provided that signaling cheater mutations are at least as likely to occur in a population as non-signaling cheater mutations.

### 2.4 The destructive or constructive nature of QS depends on cost and constitutive growth

The phase diagram in Fig. 1B demonstrates the role of both public good production cost and constitutive growth rate in determining the nature of QS outcomes. At low constitutive growth rates, public good production is essential to the survival of the population. With low λ_0_, QS regulation of public good production moves to the “mixed” region of the phase diagram at relatively low public good costs. This means that whether QS is beneficial to the population depends on the initial composition of the population. For low constitutive growth rates below λ_0_ = 0.1, all values of public good cost investigated lead to a mix of destructive and constructive cheater suppression. With larger constitutive growth rates the transition from destructiveness to constructiveness with increasing cost *c* occurs gradually with the transition to constructiveness completing at *c* = 0.1 to *c* = 0.15, depending on λ_0_. For all (c, λ_0_) pairs with *c* > 0, the cheater fixation probability is reduced in QS as compared with AO S7.

We show the ratio 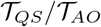 for all (n,m) pairs for two sequences of (*c*, λ_0_) pairs in Figs. S8 and S9. With fixed *c* = 0.2, the ratio 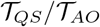 is always highest with intermediate mixtures of producers and cheaters (Fig. S8). When λ_0_ ≈ 0.1, there is a transition from the mixed region to the constructive region, where (*n, m*) pairs with few cheaters appear to be slower to change to constructiveness with increasing λ_0_. With fixed λ_0_ = 0.2, there is a gradual transition from destructiveness at low *c* to constructiveness at high *c* (Fig. S9). At moderate costs below *c* = 0.1, QS is destructive, with 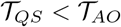 for all (*n, m*) pairs. However, once the cost reaches 0.15, QS is no longer destructive, as the mean extinction time increases with respect to AO. In the transition range from *c* = 0.1 to *c* = 0.15, the ratio 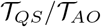 is highest when the number of producers is low, suggesting that QS is most beneficial with those initial population structures. The QS gain in mean extinction time over AO is further increased by QS at a cost of 0.2, again most drastically when *n* is small.

## 3 Discussion

Our model of competitive growth between bacterial producers and cheaters reveals a number of previously unexplored consequences of quorum sensing. When public good production allows for metabolism of otherwise inaccessible sources of nutrients, we demonstrate the important role of both the availability of alternative, public good-independent nutrients and the cost of public good production in determining the consequences of quorum sensing. With no alternative nutrients, quorum sensing by producers withholds the only source of nutrients from the whole population, which harms both producers and cheaters in terms of the total population’s mean extinction time. We identify this result as analogous to evolutionary spite, as quorum sensing reduces the absolute fitness of both producers and cheaters in terms of mean extinction time. If instead there is an alternate energy source, then regulating public good production via quorum sensing reduces the probability of cheater fixation, while in some cases increasing the mean extinction time of the population over the always on strategy and in other cases decreasing the mean extinction time, depending on the public good production cost.

The destructive region in Fig. 1 can be understood heuristically by considering that, if the cost of public good production is low, QS-mediated regulation of public good production withholds the fitness benefits of the public good while achieving minimal metabolic savings for the producers. In addition, large alternate sources of nutrition can help to offset the costs of public good production, reducing loss in relative producer fitness due to costs. On the other hand, when public good production is costly, QS-mediated production balances costs and benefits by delaying public good production until the population of producers is large enough to benefit. Thus both public good production cost and constitutive growth rate play an important role in determining the evolutionary effects of QS-based regulation.

Destructive and constructive cheater suppression is reminiscent of evolutionary spite and selfishness, respectively, as defined in previous work. However, the phenomena observed in our model are distinct from spite and selfishness for several reasons. First, the definitions of our terms are based on inter-strain relationships and the two strategies of QS and AO. This is in contrast with the Hamiltonian framework for social evolution [29, 30], where one analyzes the costs and benefits between an individual and all other individuals on the receiving end of an action. Second, by using QS instead of AO, producers harm cheaters by reducing their access to public goods when producer population density is low. However, it is unclear whether the act of withholding public goods to benefit closely related individuals can truly be called spite, in contrast to the active harm done by the costly production of bacteriocidins by *P. aeruginosa* [31], for example. Third, in our model, there is no third strain with relatively high relatedness to the producers that could indirectly benefit from cheater suppression, an accepted mechanism for the evolution of spite [23]. Finally, our study of cheater fixation probability and mean extinction time does not translate cleanly to the Hamiltonian framework [29, 30] and we can not say anything conclusive about whether the observed behavior is likely to evolve, only that it is possible in small populations. A model considering growth of small subpopulations linked by migration could provide insight into how the balance of cheater fixation and population extinction impacts the fate of the larger population, as we might expect to see in nature. This type of model could provide insight into the evolution of destructive and constructive cheater suppression and will require future work.

It would be interesting to explore the impact of mutation, migration, and horizontal gene transfer on our results, where a stable heterogeneous steady state could possibly be achieved. In addition, one could explicitly consider the role of compartmentalization through a spatial component in the model, which could also incorporate the linkage of limited dispersal and kinship [6] in terms of public goods production. Finally, the effects of heterogeneity in QS [32, 33, 34] on our results could be explored.

## 4 Materials and Methods

### 4.1 Model overview

We consider well-mixed populations of bacteria growing according to birth-death models with no mutation or migration (Fig. 1A). Public goods are modeled implicitly through their costs and benefits as reflected in the birth rates. We write the birth rates as 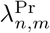 and 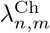 for producers and cheaters, respectively. The subscript *n* and *m* are the number of producers and cheaters present, reflecting the dependence of the rates on the population state. The death rates are 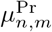 and 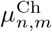 See Eqs. 1–4 for full forms of the birth and death rates.

Our model accounts for AI and public good production and degradation as well as density-dependent fitness benefits of public goods directly in the birth rates. This simplification rests on several assumptions. We assume that the timescales associated with AI and public good production and degradation are much faster than those of births and deaths. Under this assumption, the AI and public good concentrations quickly reach a steady state upon a bacterial birth or death, so that the effects of both concentrations on the birth rates is only a function of the state of the population. Similarly, the growth benefits of public goods are assumed to be a function of the current population state, namely a non-decreasing, saturating function of *n*. Another assumption is that all individuals within a given strain (producer or cheater) exhibit exactly the same birth and death rates. In reality, the stochasticity of subcellular events and diffusion of extracellular molecules leads to heterogeneous growth rates even in clonal populations [34]. Our approach simplifies analysis while providing a description of mean birth and death rates.

In order to evaluate the evolutionary success of mixed producer-cheater populations, we calculate the cheater fixation probability (*π*^Ch^), mean first passage time to cheater fixation (*τ*^Ch^), and mean population extinction time 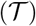 for all pairs of *n* producers and *m* cheaters where *n* + *m* ≤ *N*. We define *N* as a maximal population size large enough that the line *n* + *m* ≤ *N* effectively acts as a reflecting boundary. An appropriate choice of *N* then depends on the birth and death rates used. Solutions of all three quantities of interest come from solving backward Kolmogorov equations, as detailed in the Materials and Methods section. We validate the results from these solutions through stochastic simulations [35, 36].

### 4.2 Birth and death rates

We use birth-death models to describe the population dynamics of a bacterial colony with both public good producers and cheaters. We assume the timescales of both autoinducer and public good production and degradation are much shorter than those of population growth. Consequently, we write per capita birth and death rates solely as a function of the number of producers and cheaters in the population. We also assume that QS is sensitive to population density. While a number of other explanations for the function of QS exist, as noted in the Introduction, all of them depend to some degree on population density.

The birth and death rates in our models take into account the costs and benefits of public good production. We use λ to denote birth rates and *μ* to denote death rates, with the superscript Pr for producers and Ch for cheaters. Let *n* be the number of producers in the population and *m* be the number of cheaters. Let λ_0_ be the birth rate due to an alternative energy source that does not require proteases for bacteria to utilize, and let *μ*_0_ be the constant death rate. We assume that the per-capita death rate increases linearly with the total population size through some density-dependent mechanism such as the production of toxic byproducts. We denote the maximal growth rate due to nutrients derived from protease activity as *g* and the maximal cost of protease production, in terms of growth rate, as *c*. Parameters *K*_g_ and *K*_a_ control the number of producers for which protease-derived growth rate and protease production are half-maximal, respectively. Finally, *h*_g_ and *h*_a_ control the shape of the protease-derived growth rate and protease production rate functions. The full birth and death rates are

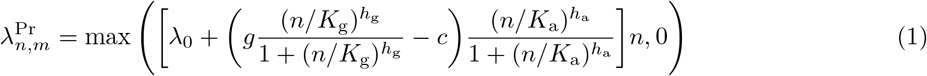

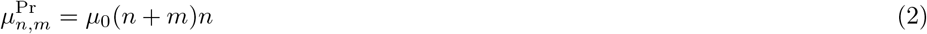

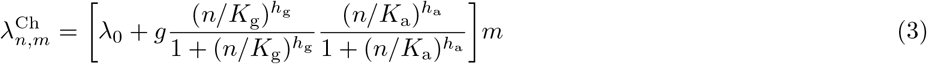

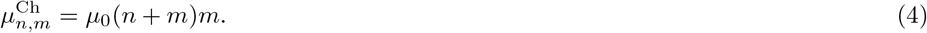

The Hill function forms of these rates are similar to those in previous work [37, 38]. The strategy taken by a particular strain in controlling public good production is controlled by the parameters *K*_a_ and *h*_a_. The parameter *K*_a_ controls when public good production is half-maximal, while *h*_a_ controls the shape of the activation curve. For an “always-on” strain, we take the limit *K*_a_ → 0 so that public good production is always maximal, independent of the producer population size. For a “no production” strain, we take *K*_a_ → ∞ so that the public good is never produced.

We set the maximum public good birth rate to *g* = 1 in all cases, so that all other birth rate parameters can be considered to be in units of maximum public good birth rate. In general we consider the parameters *g, K*_g_, *h*_g_, *c*, and λ_0_ to be properties of the public good and environment which cannot be changed by bacterial strategy. Similarly, we consider the parameter *μ*_0_ to be a joint property of the bacterial species and the environment. As we have stated, *K*_a_ and *h*_a_ constitute the choice of public good regulation strategy.

For the special case where cheaters produce autoinducer but not public goods, we modify the birth rates to

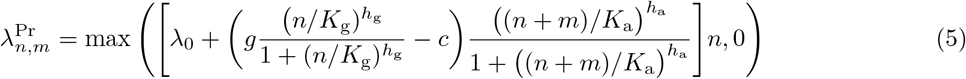

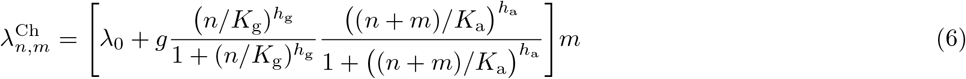

under the assumption that producers and cheaters contribute autoinducer equally. The parameter values we have used are summarized in Table 1.

#### Fixation probabilities and mean first passage times

The dynamics of the probability distribution over *n* and *m* at time *t* given *n*′ and *m*′ at an earlier time *s* (with *s* < *t*) is governed by the forward Kolmogorov equation

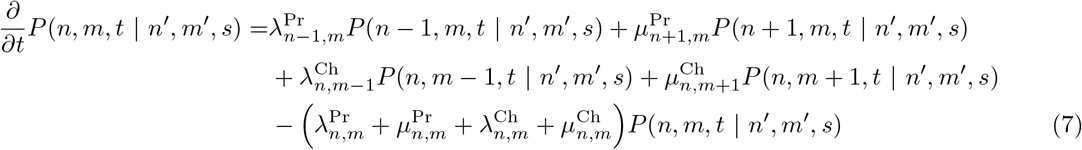

where *n, m, n*′, *m*′ ∈ {0,1, 2,… }. For birth and death rates linear in *n* and *m*, the system of coupled differential equations represented by Eq. 7 can be solved exactly using a generating function approach [39]. However, the birth and death rates required for our purposes are nonlinear, and we adopt a computational approach.

The main quantities of interest in this work are the probability of cheater fixation and the mean population extinction time. These quantities can be calculated from the backward Kolmogorov equation

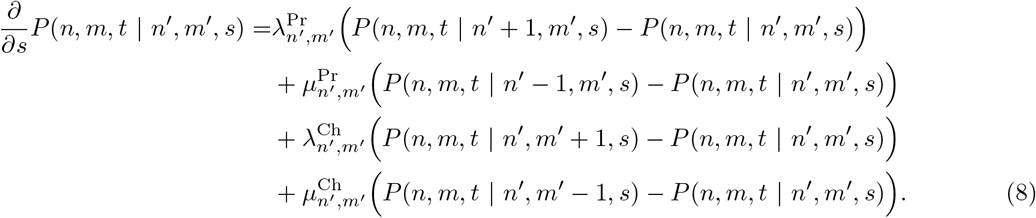

We define a maximum pure producer or cheater population size *N* and construct an (*N* + 1)^2^ × (*N* + 1)^2^ matrix describing the transition rates between all states and use it to solve for our quantities of interest. Given the birth and death rates we use, we assume that the approximation introduced by considering a finite rate matrix is reasonable when net growth rates are negative and have a large magnitude in all states where *m* = *N* or *n* = *N*. For the probability that cheaters will fix in the population, we solve [39]

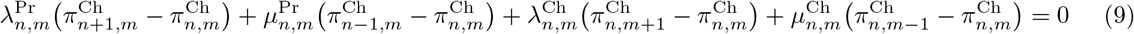

with boundary conditions

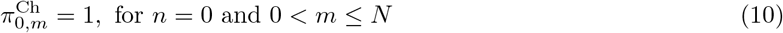

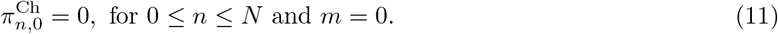

For the mean first passage time conditioned on cheater fixation, we solve

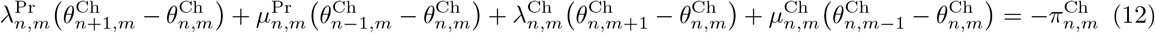

with the condition

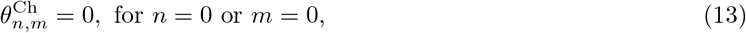

and then find the conditional mean first passage time to cheater fixation according to

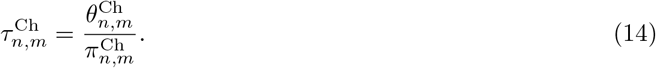

Similarly, for the mean extinction time we solve

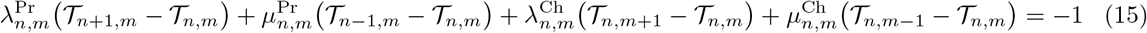

with the condition

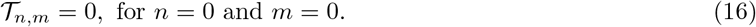

Directly solving the linear system of equations represented by Eq. 15 can lead to numerical instability when the mean extinction times are large. To account for these numerical issues we also calculated the mean extinction time using the analytical formula for pure producer or cheater populations with adjoint reflecting boundary conditions at *n* = *N* and *m* = *N* [40, 41, 42]

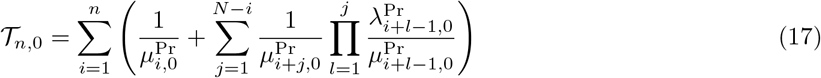

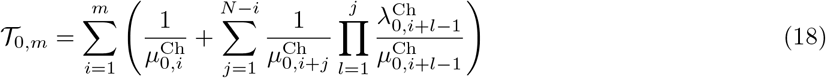

and direct calculations of fixation probabilities for each point along the pure producer and cheater axes. We use the notation 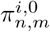 and 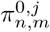 to indicate the probability that a population starting with *n* producers and *m* cheaters will have producers fix with *i* total producers and will have cheaters fix with *j* total cheaters, respectively. We then estimate the mean extinction time according to

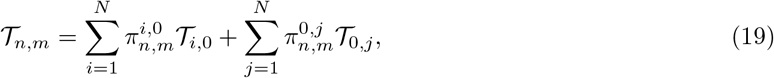

under the assumption that producers or cheaters will quickly fix followed by a much longer period with a pure population before extinction.

### 4.3 Stochastic simulations

Stochastic simulations were performed using Gillespie’s algorithm [35, 36] implemented in the Gillespy2 Python package [43]. Each simulation was repeated 100 times in order to estimate fixation probabilities and mean first passage times.

### 4.4 General methods

The figures in this article were created using Inkscape 0.92 and Matplotlib 3.1.1 [44] in a Python 3.7 Jupyter Notebook [45]. All fixation probabilities and mean first passage times were calculated using the linear system solver in the Python NumPy package [46] and the sparse matrix module of the SciPy package [47].

## Acknowledgments

A.S.M. and A.W.E. were partially supported by the United States Defense Advanced Research Projects Agency RadioBio program under grant number HR001117C0125, and by a Discovery grant from the Natural Sciences and Engineering Research Council of Canada. P.J.T. was partially supported by National Science Foundation grant DMS-2052109, and by the Oberlin College Libraries.

## A Supplementary Figures and Tables

**Figure S1:**
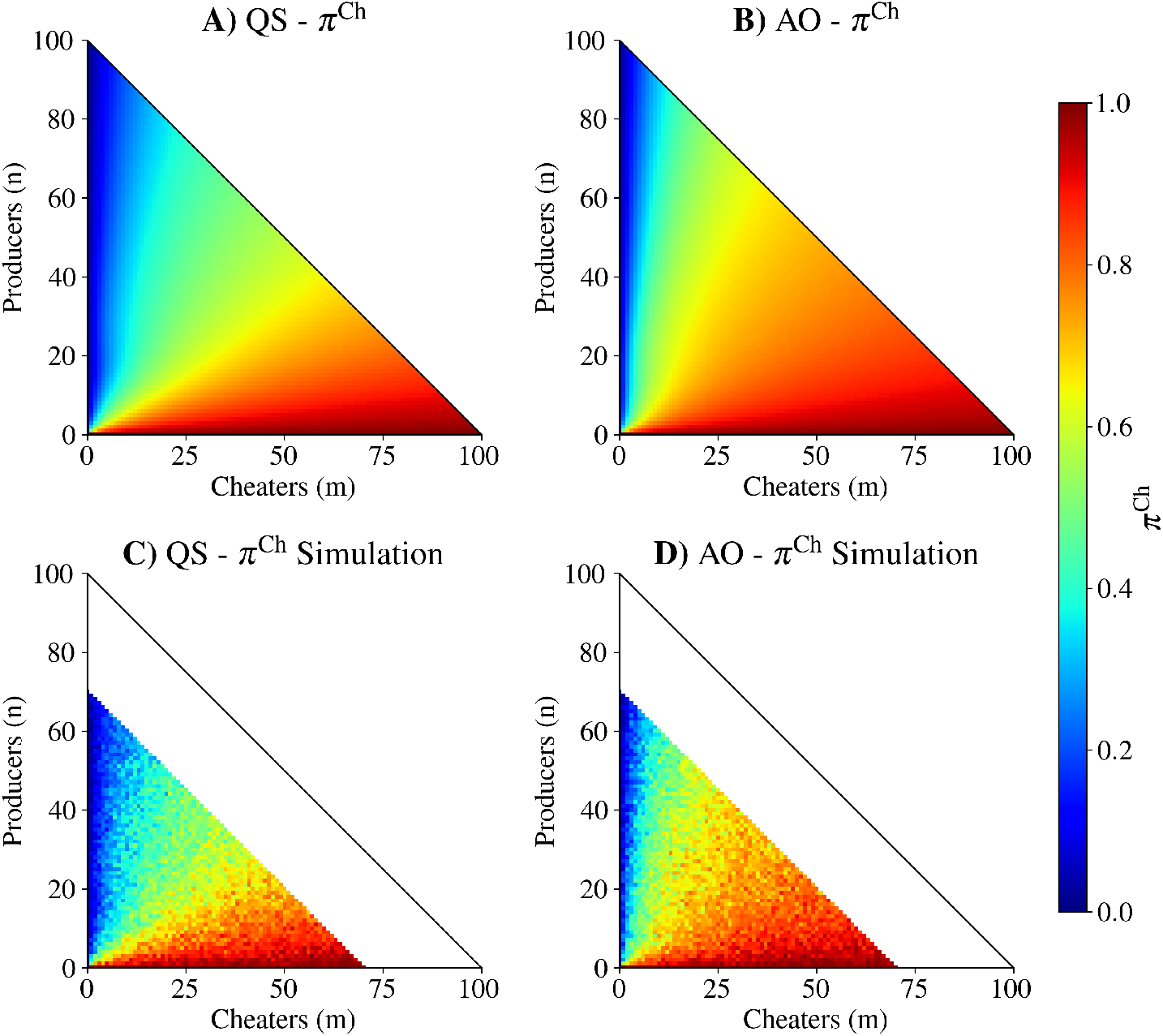
The cheater fixation probabilities from Fig 2 agree with simulation results. The first row shows the cheater fixation probabilities for A) quorum sensing (QS) and B) always on (AO) strategies, directly reproduced from Fig. 2. The second row shows cheater fixation probabilities calculated as a mean from 100 independent simulations for C) QS and D) AO strategies. Points above *n* + *m* = 70 (above the zero-net growth contour) were not calculated. See Table 2 for parameter values.

**Figure S2:**
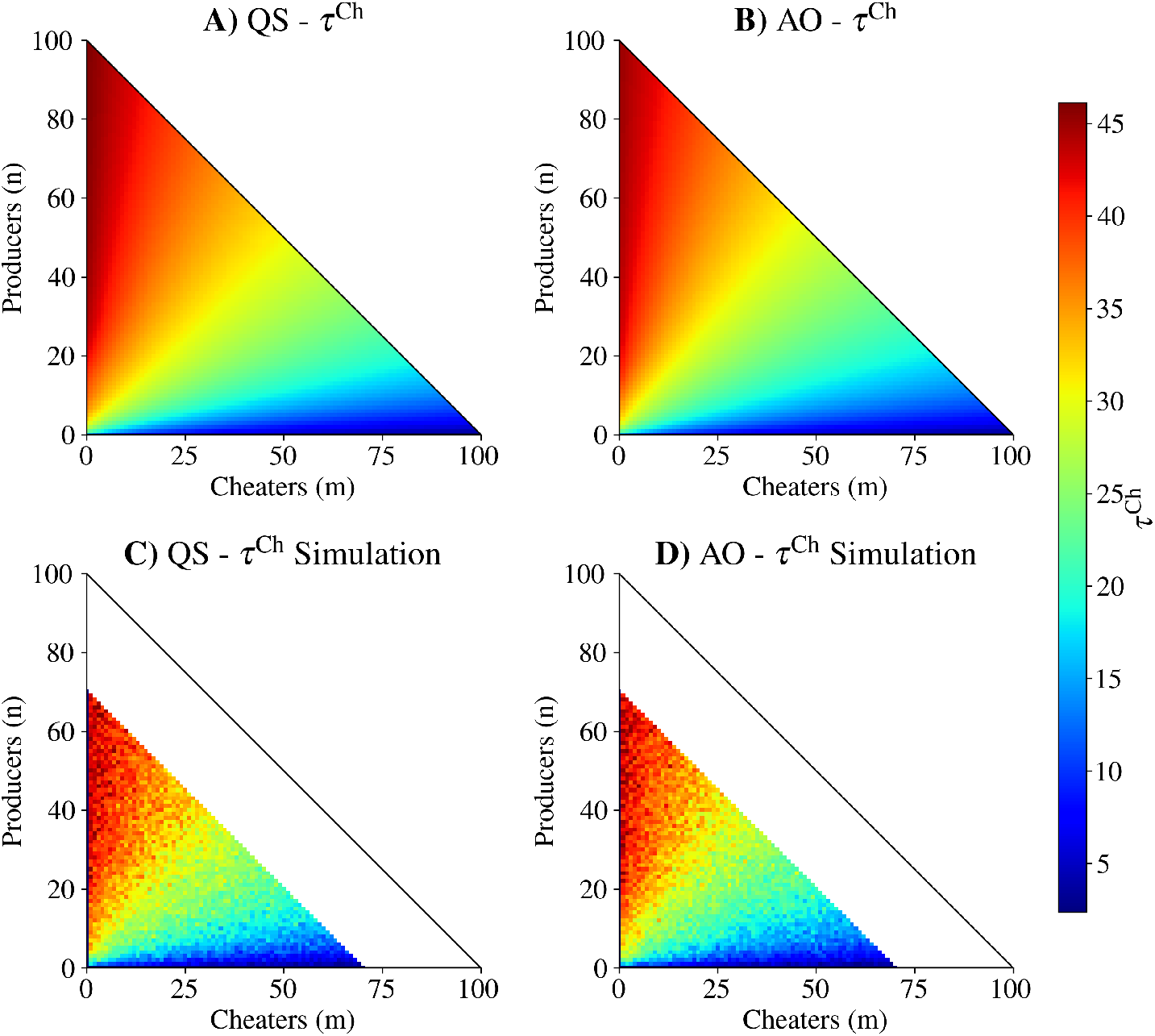
The cheater mean first passage time to fixation from Fig 2 agree with simulation results. The first row shows the cheater fixation mean first passage times for A) quorum sensing (QS) and B) always on (AO) strategies, directly reproduced from Fig. 2. The second row shows cheater fixation mean first passage times calculated as a mean from 100 independent simulations for C) QS and D) AO strategies. Points above *n* + *m* = 70 (above the zero-net growth contour) were not calculated. See Table 2 for parameter values.

**Figure S3:**
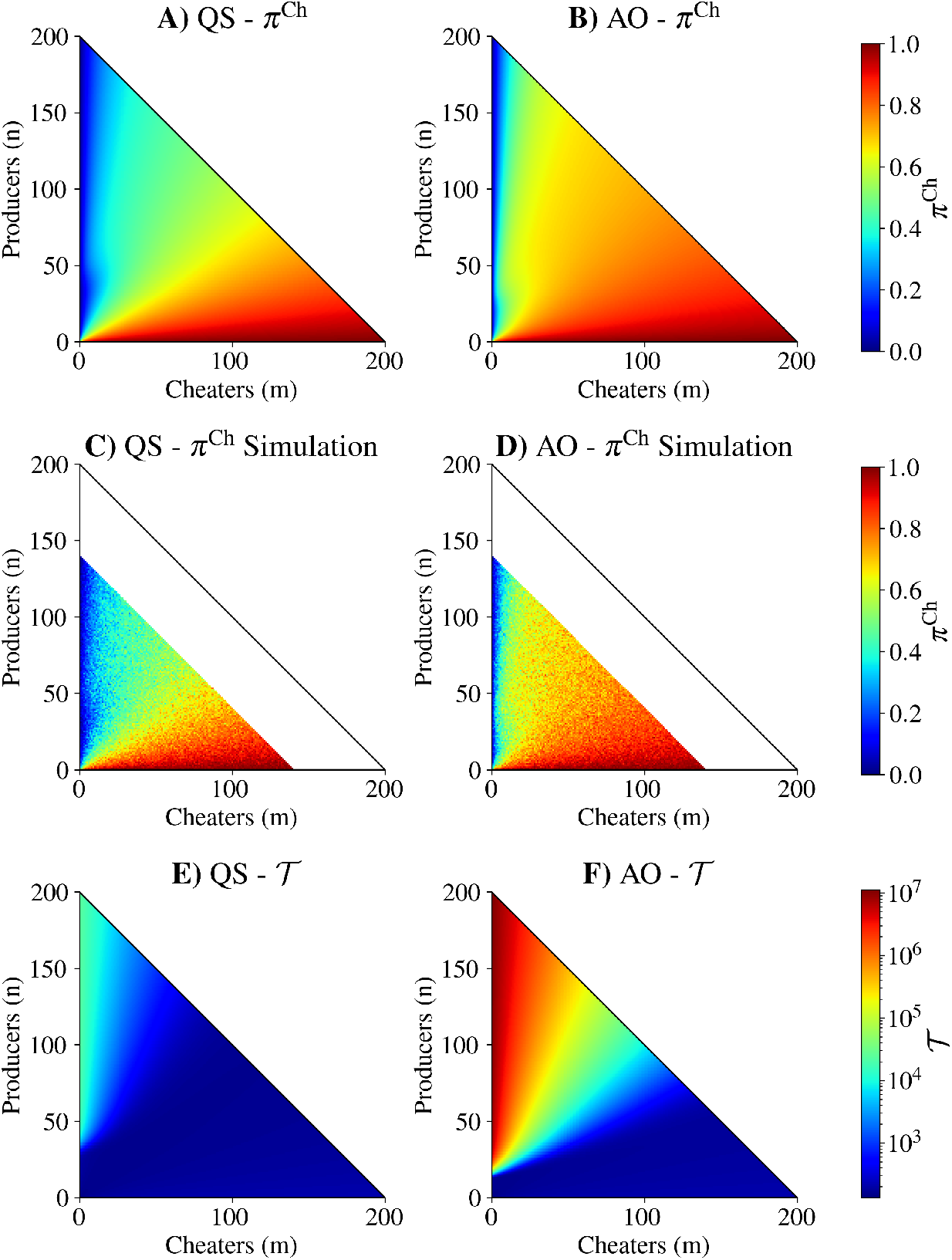
The qualitative results from Fig. 2 are preserved with increased carrying capacity, with no constitutive growth (λ_0_). The first row depicts cheater fixation probability from an initial population structure of *n* producers and *m* cheaters for A) quorum sensing (QS) and B) always on (AO) strategies. The second row depicts cheater fixation probabilities calculated as a mean of 100 independent simulations for C) QS and D) AO strategies. Points above *n* + *m* = 140 (above the zero-net growth contour) were not calculated. The third row depicts mean extinction time from initial population structure for E) QS and F) AO strategies, calculated according to Eq. 19. As with the results in Fig. 2, QS decreases cheater fixation probability but also decreases mean extinction time as compared with AO. See Table 2 for parameter values.

**Figure S4:**
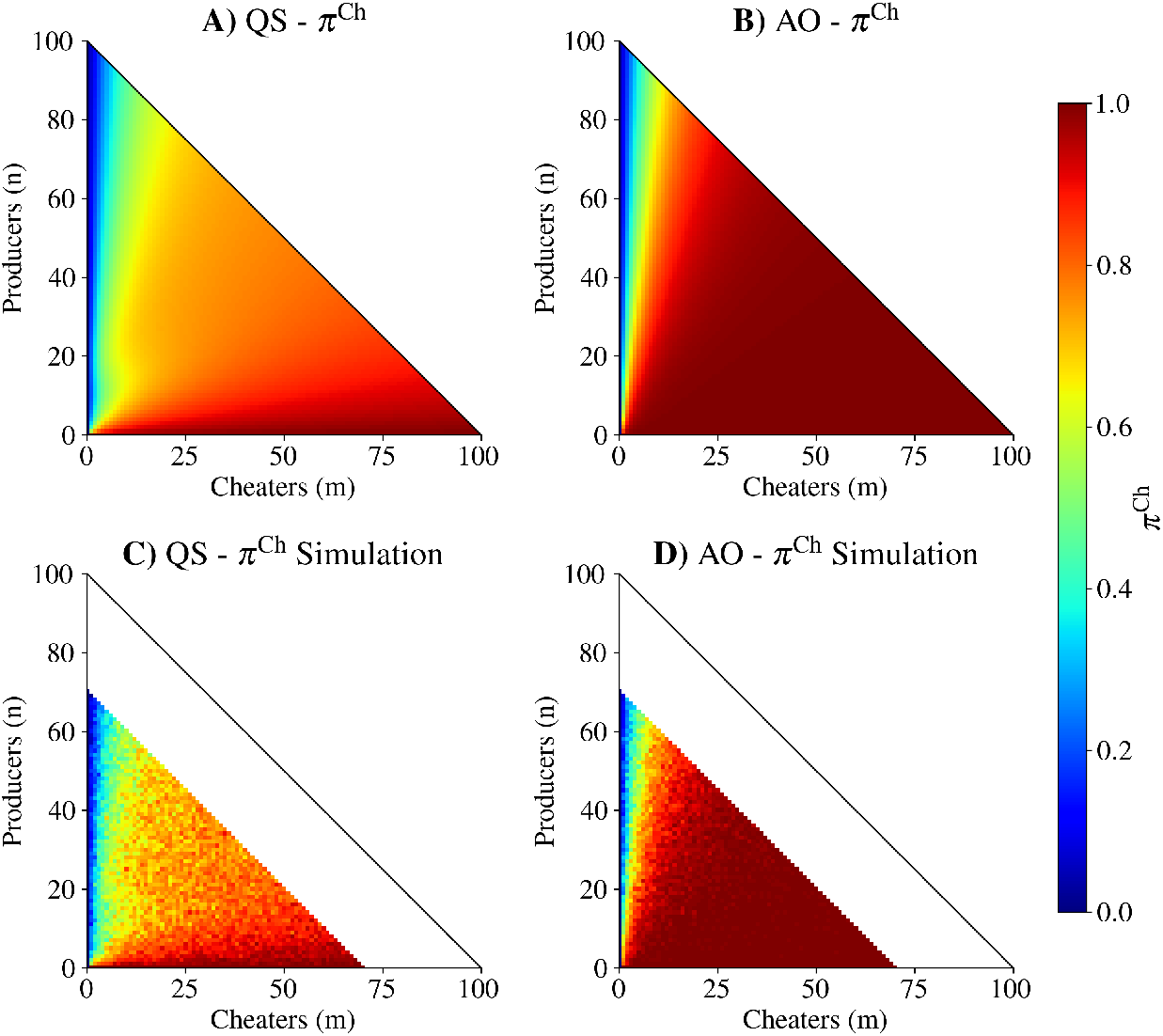
The cheater fixation probabilities from Fig 3 agree with simulation results. The first row shows the cheater fixation probabilities for A) quorum sensing (QS) and B) always on (AO) strategies, directly reproduced from Fig. 3. The second row shows cheater fixation probabilities calculated as a mean from 100 independent simulations for C) QS and D) AO strategies. Points above *n* + *m* = 70 (above the zero-net growth contour) were not calculated. See Table 2 for parameter values.

**Figure S5:**
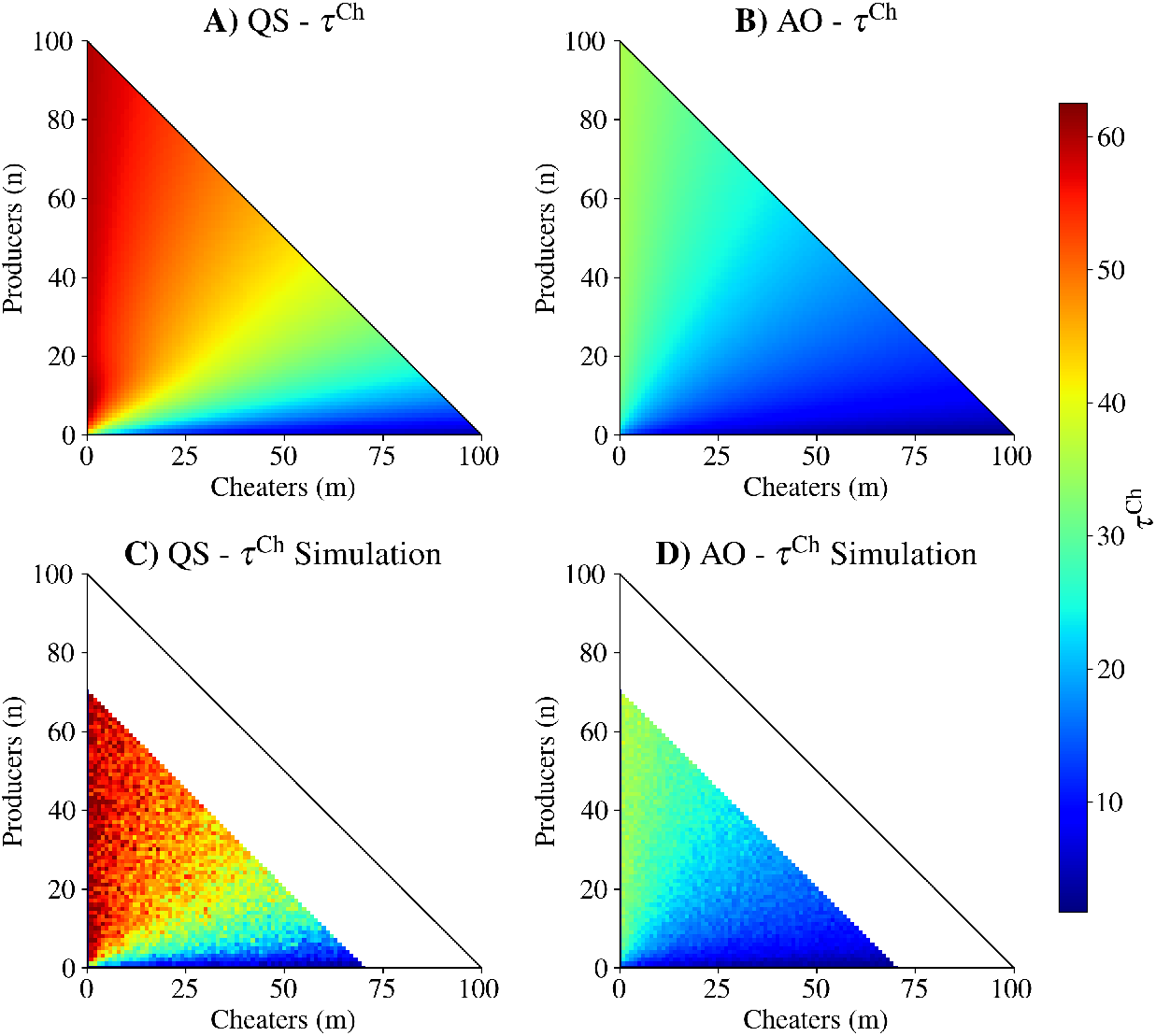
The cheater mean first passage time to fixation from Fig 3 agree with simulation results. The first row shows the cheater fixation mean first passage times for A) quorum sensing (QS) and B) always on (AO) strategies, directly reproduced from Fig. 3. The second row shows cheater fixation mean first passage times calculated as a mean from 100 independent simulations for C) QS and D) AO strategies. Points above *n* + *m* = 70 (above the zero-net growth contour) were not calculated. See Table 2 for parameter values.

**Figure S6:**
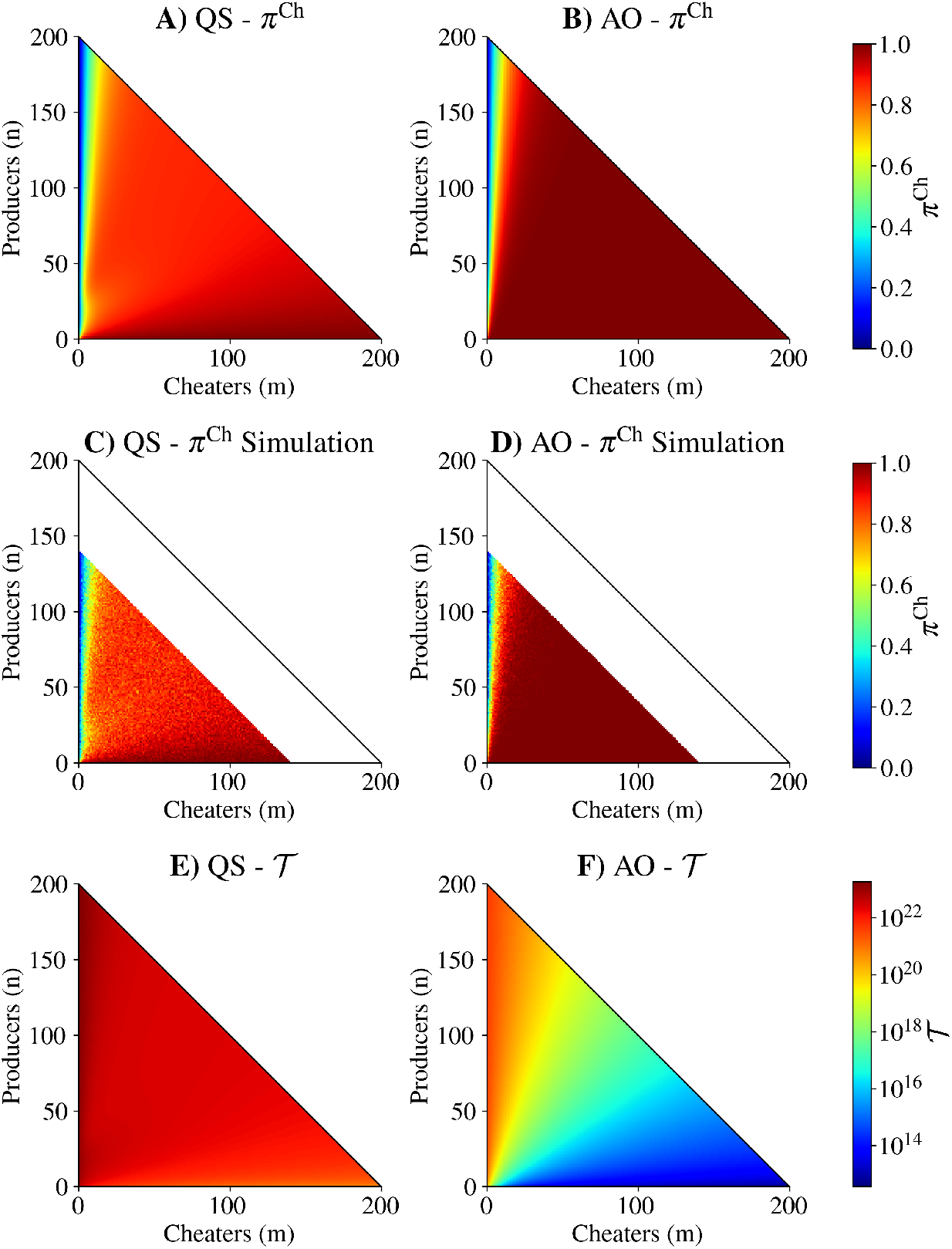
The qualitative results from Fig. 3 are preserved with increased carrying capacity, with constitutive growth (λ_0_ = 0.2). The first row depicts cheater fixation probability from an initial population structure of *n* producers and *m* cheaters for A) quorum sensing (QS) and B) always on (AO) strategies. The second row depicts cheater fixation probabilities calculated as a mean of 100 independent simulations for C) QS and D) AO strategies. Points above *n* + *m* = 140 (above the zero-net growth contour) were not calculated. The third row depicts mean extinction time from initial population structure for E) QS and F) AO strategies, calculated according to Eq. 19. As with the results in Fig. 3, QS decreases cheater fixation probability while also increasing mean extinction time as compared with AO. See Table 2 for parameter values.

**Figure S7:**
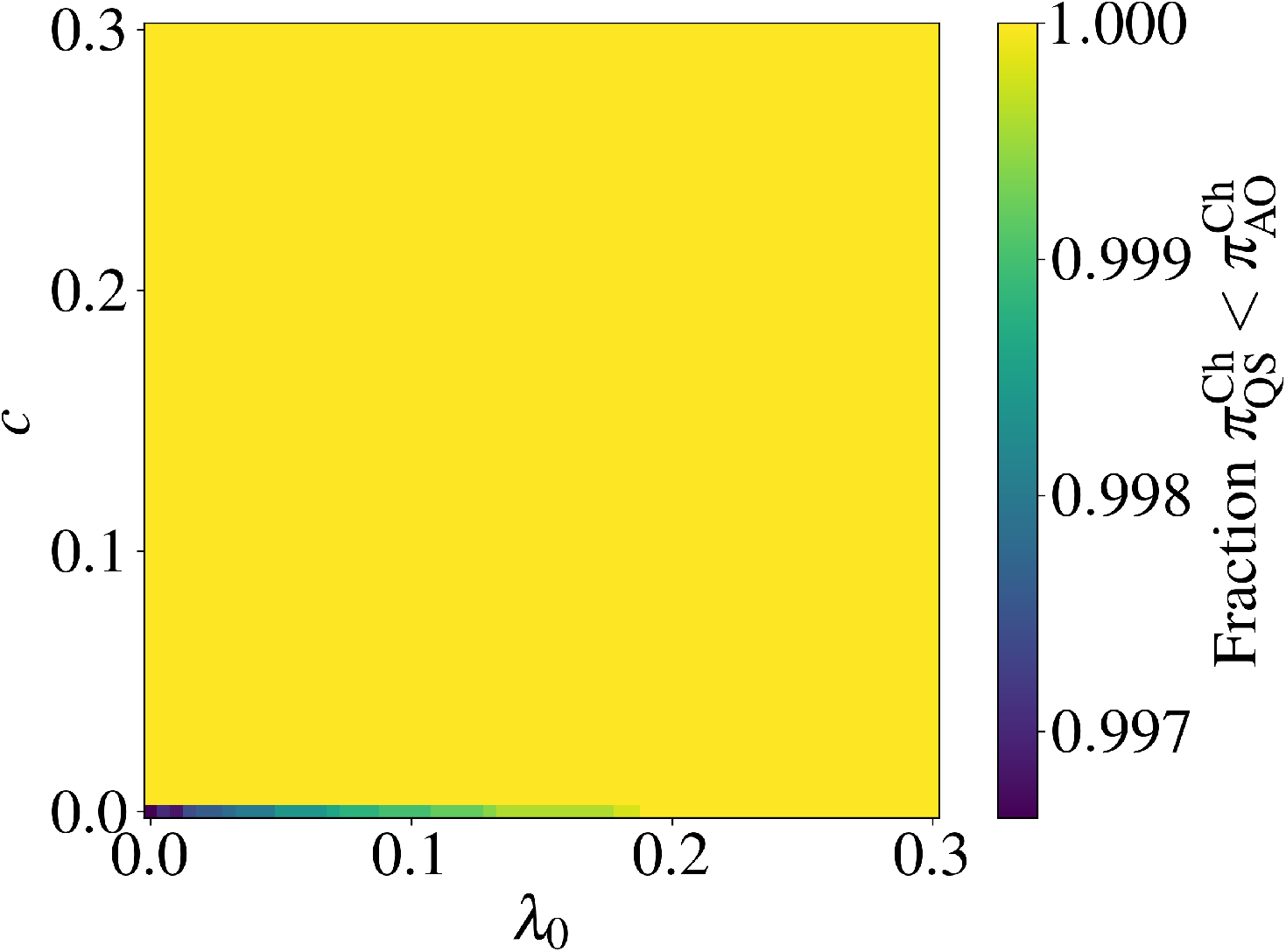
Phase diagram describing the fitness gains of QS over AO as a function of constitutive growth rate (λ_0_) and public good production cost (c). For each pair (λ_0_, *c*) we calculated the cheater fixation probability for all (*n, m*) pairs satisfying *n* ≥ 0, *m* ≥ 0, and *n* + *m* ≤ 100 with the QS and AO strategies. The reported number is the fraction of these (*n, m*) pairs where the cheater fixation probability for QS is less than for AO 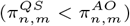. For all (λ_0_, *c*) pairs except for when *c* = 0, cheater fixation probability is reduced by QS for all initial population compositions. When *c* = 0, there are a small number of initial compositions where 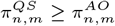 which decreases as λ_0_ increases. See Table 2 for parameter values.

**Figure S8:**
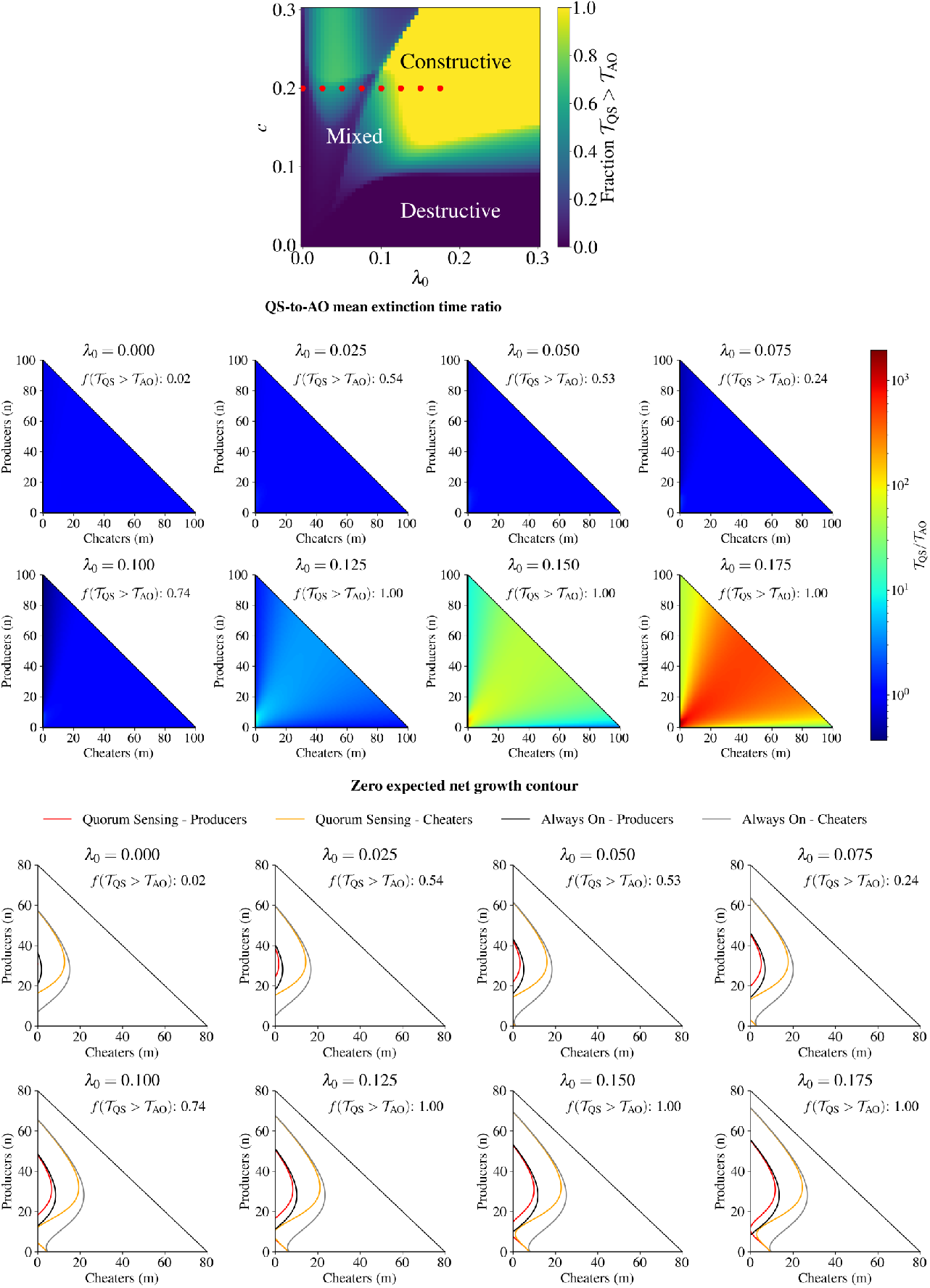
Detailed analysis of the phase diagram from Fig. 1 with *c* = 0.2 fixed and with different λ_0_ values. Top row: the red dots shown over the phase diagram indicate the (*c*, λ_0_) pairs that we examine in detail. Middle row: the ratio of QS mean extinction time to AO mean extinction time 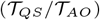 with the indicated values of λ_0_. The fraction of (*n, m*) pairs where 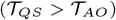 is indicated as 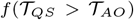. Bottom row: the zero expected net growth contour contours for QS producers (red), QS cheaters (orange), AO producers (black), and AO cheaters (grey). For λ_0_ = 0 the two QS zero expected net growth contour contours are identical and the two AO zero expected net growth contour contours are also identical. As λ_0_ is increased, the AO producer contour approaches the producer axis faster than the QS producer contour. See Table 2 for parameter values.

**Figure S9:**
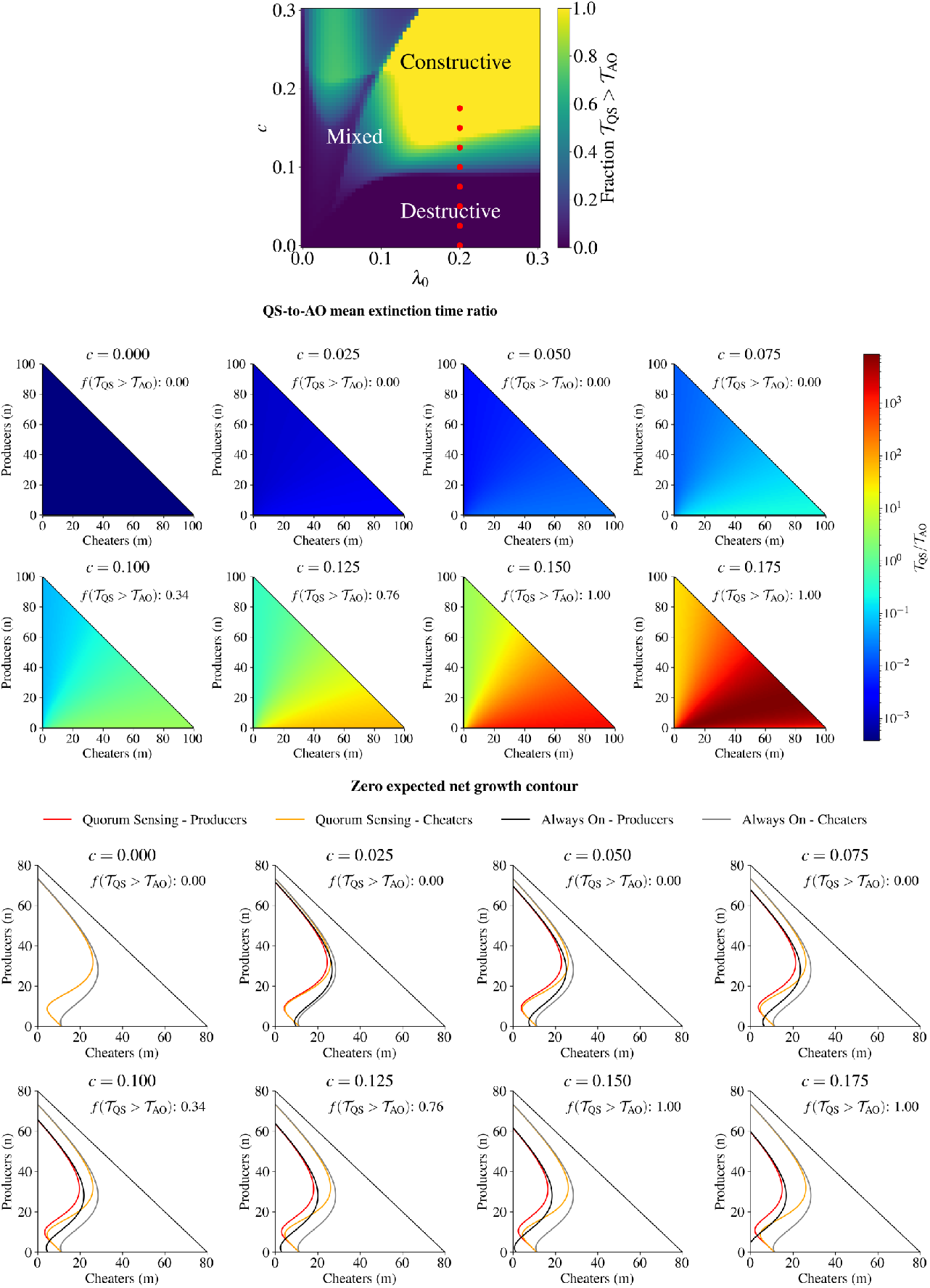
Detailed analysis of the phase diagram from Fig. 1 with λ_0_ = 0.2 fixed and with different *c* values. Top row: the red dots shown over the phase diagram indicate the (*c*, λ_0_) pairs that we examine in detail. Middle row: the ratio of QS mean extinction time to AO mean extinction time 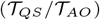 with the indicated values of *c*. The fraction of (*n, m*) pairs where 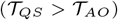 is indicated as 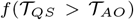. Bottom row: the zero expected net growth contour contours for QS producers (red), QS cheaters (orange), AO producers (black), and AO cheaters (grey). At λ_0_ = 0.075 the diagonal zero expected net growth contour contour near the origin appears for QS, and moves further from the origin as λ_0_ increases from there. See Table 2 for parameter values.

